# Chemotherapy weakly contributes to predicted neoantigen expression in ovarian cancer

**DOI:** 10.1101/090134

**Authors:** Timothy O’Donnell, Elizabeth L. Christie, Arun Ahuja, Jacqueline Buros, B. Arman Aksoy, David D. L. Bowtell, Alexandra Snyder, Jeff Hammerbacher

## Abstract

**Background:** Patients with highly mutated tumors, such as melanoma or smoking-related lung cancer, have higher rates of response to immune checkpoint blockade therapy, perhaps due to increased neoantigen expression. Many chemotherapies including platinum compounds are known to be mutagenic, but the impact of standard treatment protocols on mutational burden and resulting neoantigen expression in most human cancers is unknown.

**Methods:** We sought to quantify the effect of chemotherapy treatment on computationally predicted neoantigen expression for 12 high grade serous ovarian carcinoma (HGSC) patients with pre- and post-chemotherapy samples collected in the Australian Ovarian Cancer Study. We additionally analyzed 16 patients from the cohort with post-treatment samples only, including five primary surgical samples exposed to neoadjuvant chemotherapy. Our approach integrates tumor whole genome and RNA sequencing with class I MHC binding prediction and mutational signatures of chemotherapy exposure extracted from two preclinical studies.

**Results:** The mutational signatures for cisplatin and cyclophosphamide identified in a preclinical model had significant but inexact associations with the relevant exposure in the clinical samples. In an analysis stratified by tissue type (solid tumor or ascites), relapse samples collected after chemotherapy harbored a median of 90% more expressed neoantigens than untreated primary samples, a figure that combines the effects of chemotherapy and other mutagenic processes operative during relapse. Neoadjuvant-treated primary samples showed no detectable increase over untreated samples. The contribution from chemotherapy-associated signatures was small, accounting for a mean of 5% (range 0–16) of the expressed neoantigen burden in relapse samples. In both treated and untreated samples, most neoantigens were attributed to COSMIC *Signature (3)*, associated with BRCA disruption, *Signature (1)*, associated with a slow mutagenic process active in healthy tissue, and *Signature (8)*, of unknown etiology.

**Conclusion:** Relapsed HGSC tumors harbor nearly double the predicted expressed neoantigen burden of primary samples, but mutations associated with chemotherapy signatures account for only a small part of this increase. The mutagenic processes responsible for most neoantigens are similar between primary and relapse samples. Our analyses are based on mutations detectable from whole genome sequencing of bulk samples and do not account for neoantigens present in small populations of cells.

## Background

Many chemotherapies including platinum compounds [1], cyclophosphamide [2], and etoposide [3] exert their effect through DNA damage, and recent studies have found evidence for chemotherapy-induced mutations in post-treatment acute myeloid leukaemia [4], glioma [5], and esophageal adenocarcinoma [6]. Successful development of immune checkpoint-mediated therapy[7] has focused attention on the importance of T cell responses to somatic mutations in coding genes that generate neoantigens [8]. Studies based on bulk-sequencing of tumor samples followed by computational peptide-class I MHC affinity prediction [9] have suggested that tumors with more mutations and predicted mutant MHC I peptide ligands are more likely to respond to checkpoint blockade immunotherapy [10] [11]. Ovarian cancers fall into an intermediate group of solid tumors in terms of mutational load present in pre-treatment surgical samples[12]. However, the effect of standard chemotherapy regimes on tumor mutation burden and resulting neoantigen expression in ovarian cancer is poorly understood.

Investigators associated with the Australian Ovarian Cancer Study (AOCS) performed whole genome and RNA sequencing of 79 pre-treatment and 35 post-treatment cancer samples from 92 HGSC patients, including 12 patients with both pre- and post-treatment samples [13]. The samples were obtained from solid tissue resections, autopsies, and ascites drained to relieve abdominal distension. Treatment regimes varied but primary treatment always included platinum-based chemotherapy. In their analysis, Patch et al. reported that post-treatment samples harbored more somatic mutations than pre-treatment samples and exhibited evidence of chemotherapy-associated mutations. Here we extend these results by quantifying the mutations and predicted neoantigens attributable to chemotherapy-associated mutational signatures. We find that, while neoantigen expression increases after treatment and relapse, only a small part of the increase is due to mutations associated with chemotherapy signatures.

## Methods

### Clinical sample information

We grouped the AOCS samples into three sets — “primary/untreated,” “primary/treated,” and “relapse/treated” — according to collection time point and chemotherapy exposure. The primary/untreated group consists of 75 primary debulking surgical samples and 4 samples of drained ascites. The primary/treated group consists of 5 primary debulking surgical samples obtained from patients pretreated with chemotherapy prior to surgery (neoadjuvant chemotherapy). The relapse/treated group consists of 24 relapse or recurrence ascites samples, 5 metastatic samples obtained in autopsies of two patients, and 1 solid tissue relapse surgical sample, all of which were obtained after prior exposure to one or more lines of chemotherapy. In summary, these groupings yield 79 primary/untreated samples, 5 primary/treated samples, and 30 relapse/treated samples. Sample and clinical information including chemotherapy treatments is listed in Additional File 1.

Independent of treatment, ascites samples trend toward more detected mutations, perhaps due to increased intermixing of clones. We therefore stratified by tissue type (solid tumor or ascites) when comparing the mutation and neoantigen burdens of pre- and post-treatment samples.

### Mutation calls

We analyzed the mutation calls published by Patch et al. [13] (Additional File 2). DNA and RNA sequencing reads were downloaded from the European Genome-phenome Archive under accession EGAD00001000877. Adjacent SNVs from the same patient were combined to form multinucleotide variants (MNVs).

We considered a mutation to be present in a sample if it was called for the patient and more than 5 percent of the overlapping reads and at least 6 reads total supported the alternate allele. We considered a mutation to be expressed if there were 3 or more RNA reads supporting the alternate allele. In the analysis of paired pre- and post-treatment samples from the same donors, we defined a mutation as unique to the post-treatment sample if the pre-treatment sample contained greater than 30 reads coverage and no variant reads at the site.

### Variant annotation, HLA typing, and MHC binding prediction

The most disruptive effect (in terms of amino acid sequence) of each protein-changing variant was predicted using Varcode [15]. For insertions or deletions (indels) that were predicted to disrupt the reading frame, all downstream peptides potentially generated up to a stop codon were considered.

HLA typing was performed using a consensus of seq2HLA [16] and OptiType [17] across the samples for each patient (Additional File 3).

Class I MHC binding predictions were performed for peptides of length 8–11 using NetMHCpan 2.8 [18] with default arguments (predicted neoantigens are listed in Additional File 2).

### Mutational signatures

The use of mutational signatures is necessary because it is not possible to distinguish chemotherapy-induced mutations from temporal effects when comparing primary and relapse samples by mutation count alone. A mutational signature ascribes a probability to each of the 96 possible single-nucleotide variants, where a variant is defined by its reference base pair, alternate base pair, and base pairs immediately adjacent to the mutation. Signatures have been associated with exposure to particular mutagens, age related DNA changes, and disruption of DNA damage repair pathways due to somatic mutations or germline risk variants in melanoma, breast, lung and other cancers [19], and provide a means of identifying the contribution that chemotherapy may make to the mutations seen in post-treatment samples. For example, the chemotherapy temozolomide has been shown to induce mutations consisting predominantly of *C* → *T* (equivalently, *G* → *A*) transitions at CpC and CpT dinucleotides [5]. To perform deconvolution, the single nucleotide variants (SNVs) observed in a sample are tabulated by trinucleotide context, and a combination of signatures, each corresponding to a mutagenic process, is found that best explains the observed counts. Mutational signatures may be discovered *de novo* from large cancer sequencing projects but for smaller studies it is preferable to deconvolve using known signatures [20].

The Catalogue Of Somatic Mutations In Cancer (COSMIC) Signature Resource curates 30 signatures discovered in a pan-cancer analysis of untreated primary tissue samples. While signatures for exposure to the chemotherapies used in ovarian cancer have not been established from human studies, two recent reports provide data on mutations detected in cisplatin-exposed *C. Elegans* [21] and a *G. Gallus* cell line exposed to several chemotherapies including cisplatin, chyclophosphamide, and etoposide [22]. From the SNVs identified in these studies, we defined two signatures for cisplatin, a signature for cyclophosphamide, and a signature for etoposide (Figures S1 and S2). As both studies sequenced replicates of chemotherapy-treated and untreated (control) samples, identifying a mutational signature associated with treatment required splitting the mutations observed in the treated group into background and treatment effects. We did this using a Bayesian model for each study and chemotherapy drug separately.

Let *C*_*i,j*_ be the number of mutations observed in experiment *i* for mutational trinucletoide context 0 ≤ *j* < 96. Let *t*_*i*_ ∈ {0,1} be 1 if the treatment was administered in experiment *i* and 0 if it was a control.

We estimate the number of mutations in each context arising due to background (non-treatment) processes *B*_*j*_ and the number due to treatment *T*_*j*_ according to the model:

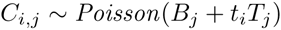

We fit this model using Stan [23] with a uniform (improper) prior on the entries of *B* and *T*. The treatment-associated mutational signature *N* was calculated from a point estimate of *T* as:

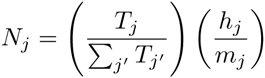

where *h*_*j*_ and *m*_*j*_ are the number of times the reference trinucleotide *j* occurs in the human and preclinical model (*C. Elegans* or *G. Gallus*) genomes, respectively.

Signature deconvolution was performed with the deconstructSigs[20] package using the 30 mutational signatures curated by COSMIC [24] extended to include the putative chemotherapy-associated signatures (Additional Files 4 and 5). When establishing whether a signature was detected in a sample, we applied the 6% cutoff recommended by the authors of the deconstructSigs package. Signatures assigned weights less than this threshold in a sample were considered undetected.

To estimate the number of SNVs and neoantigens generated by a signature, for each mutation in the sample we calculated the posterior probability that the signature generated the mutation, as described below. The sum of these probabilities gives the expected number of SNVs attributable to each signature. For neoantigens, we weighted the terms of this sum by the number of neoantigens generated by each mutation.

Suppose a mutation occurs in context *j* and sample *i*. We calculate Pr[*s* | *j*], the probability that signature *s* gave rise to this mutation, using Bayes’ rule:

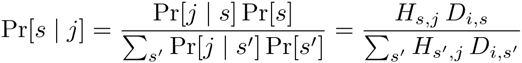

where *D*_*i,s*_ gives the contribution of signature *s* to sample *i* and *H*_*s,j*_ is the weight for signature *s* on mutational context *j*. For treated samples with a pre-treatment sample available from the same patient, we deconvolved signatures for both the full set of mutations and for the mutations detected only after treatment. When calculating Pr[*s* | *j*] for these samples, for each mutation we selected the appropriate deconvolution matrix *D*_*i*,*s*_ based on whether the mutation was unique to the post-treatment sample.

## Results

### Cisplatin and cyclophosphamide mutational signatures correlate with clinical treatment

We identified mutational signatures for cisplatin, cyclophosphamide, and etoposide from the *G. Gallus* cell line data (Figure S1), as well as a second cisplatin signature from experiments in *C. Elegans* (Figure S2). The two cisplatin signatures were not identical. Both signatures placed most probability mass on *C → A* mutations, but differed in preference for the nucleotides adjacent to the mutation. The *G. Gallus* signature was relatively indifferent to the 5’ base and favored a 3’ cytosine, whereas the *C. Elegans* signature was specific for a 5’ cytosine and a 3’ pyrmidine. The *G. Gallus* cisplatin signature was closest in cosine distance to COSMIC *Signature (24) Aflatoxin*, *Signature (4) Smoking*, and *Signature (29) Chewing tobacco*, all associated with guanine adducts. The *C. Elegans* cisplatin signature was similar to *Signature (4) Smoking*, *Signature (20) Mismatch repair*, and *Signature (14) Unknown*. The *G. Gallus* cyclophosphamide signature favored *T → A* and *C → T* mutations and was most similar to COSMIC Signatures *(25)*, *(8)*, and *(5)*, all of unknown etiology. The *G. Gallus* etoposide signature distributed probability mass nearly uniformly across mutation contexts and was most similar to COSMIC *Signature (5) Unknown*, *Signature (3) BRCA*, and *Signature (16) Unknown*. Overall, the chemotherapy signatures were no closer to any COSMIC signatures than the two most similar COSMIC signatures (*Signature (12) Unknown* and *Signature (26) Mismatch repair*) are to each other, suggesting that deconvolution could in principle distinguish their contributions.

We performed signature deconvolution on each sample’s SNVs (top and middle of Figures S3 and ??). Detection of the cyclophosphamide signature at the 6% threshold was associated with clinical cyclophosphamide treatment (Bonferroni-corrected Fischer’s exact test *p* = 0.004), occurring in 4/10 samples taken after cyclophosphamide treatment, 2/79 pre-treatment samples, and 2/25 samples exposed to chemotherapies other than cyclophosphamide. In contrast, the two cisplatin signatures were found in no samples, and the etoposide signature was found only in four pre-treatment samples.

For better sensitivity, we next focused on the 14 relapse/treated samples from the 12 patients with both pre- and post-treatment samples. For each patient, we extracted the mutations that had evidence exclusively in the treated samples. Of 206,766 SNVs in the post-treatment samples for these patients, 93,986 (45%) satisfied our filter and were subjected to signature deconvolution (Figure 1, bottom of Figures S3 and ??). Within this subgroup, the *G. gallus* cisplatin signature was identified only in the two samples taken after cisplatin therapy, a significant association (*p* = 0.04). The *C. Elegans* cisplatin signature was detected in no samples, and the cyclophosphamide signature was detected in 3/6 cyclophosphamide-treated samples, but, unexpectedly, also in 6/8 non-cyclophosphamide-treated samples. These included the two post-treatment samples in which the signature was detected in the earlier analysis plus four additional samples. COSMIC *Signature (3) BRCA* and *Signature (8) Unknown etiology* were detected in 14/14 and 9/14 post-treatment samples, respectively, but *Signature (1) Age* was absent, consistent with its association with a slow mutagenic process operative before oncogenesis.

**Figure 1:**
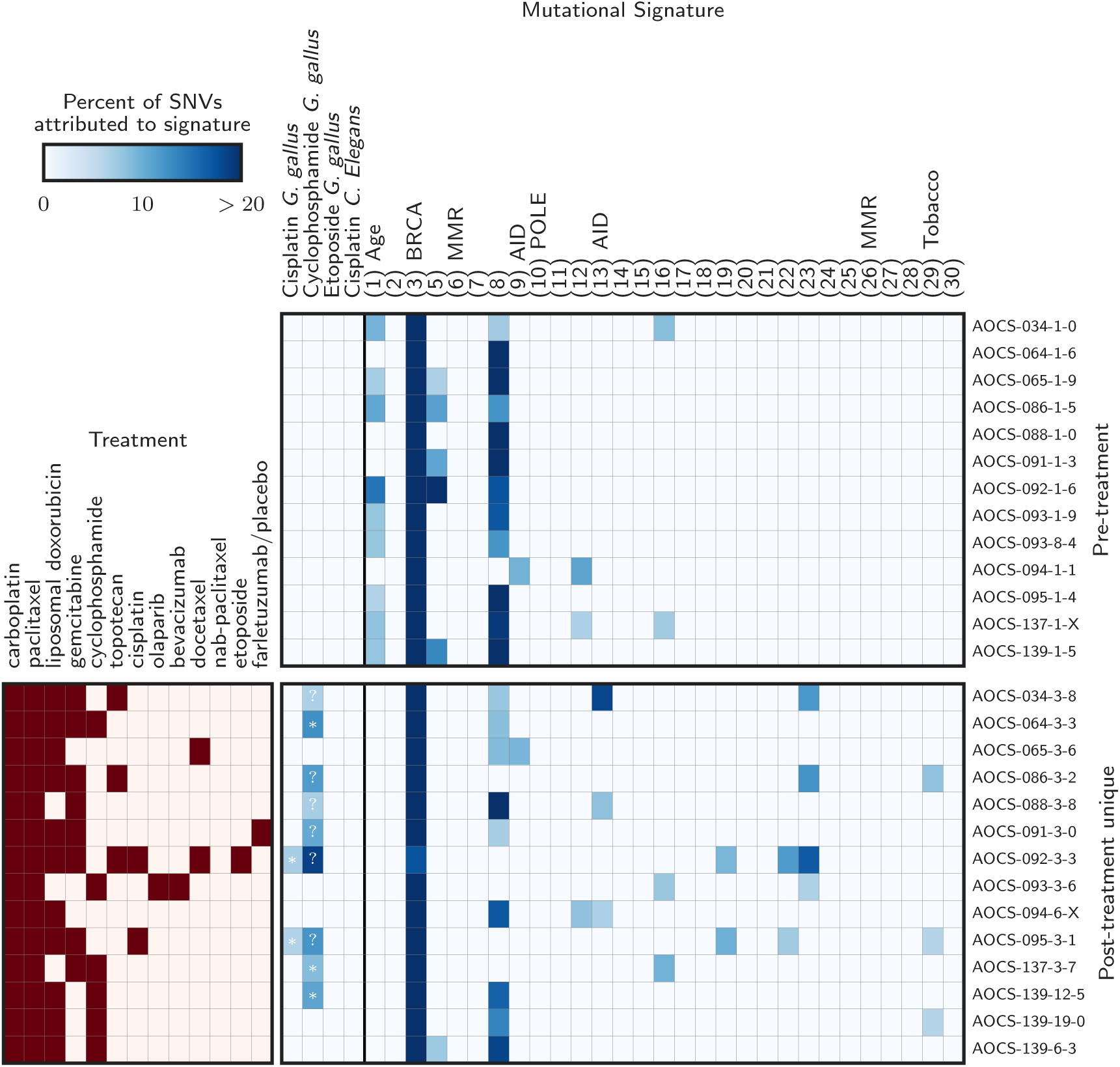
Detected mutational signatures for donor-matched primary/untreated and relapse/treated samples. *(Top)* Signatures detected in the pre-treatment samples. The first four signatures were extracted from reports of a *G. gallus* cell line and *C. Elegans* after exposure to chemotherapy, and the rest are COSMIC curated signatures. COSMIC signature numbers are shown in parentheses, and the associated mutagenic process is indicated when known. Signatures not shown were undetected in these samples. *(Bottom)* Clinical treatments and detected signatures for the mutations unique to the post-treatment samples (those with no evidence in the matched pre-treatment sample). Cases where a chemotherapy signature is detected are annotated with a (*) if the patient received the associated drug and a (?) otherwise.

In summary, the mutational signatures for cisplatin and cyclophosphamide extracted from experiments of a *G. Gallus* cell line showed significant but inexact associations with clinical chemotherapy exposure.

### Neoantigen burden increases at relapse

Across the cohort, we identified 17,689 mutated peptides predicted to bind autologous MHC class I with affinity 500nm or tighter [25]. All but 21 (0.12%) of these predicted neoantigens were private to a single patient (shared neoantigens are listed in Additional File 6).

Relapse/treated samples showed more expressed neoantigens than primary/untreated samples. Solid tissue relapse samples harbored a median of 81% (bootstrap 95% CI 40–123) more mutations, 124% (58– 191) more neoantigens, and 90% (40–142) more expressed neoantigens than primary/untreated solid tissue samples (Figure 2), all significant increases (Mann-Whitney *p <* 0.004 for each of the three tests). A similar trend was observed for ascites samples. Relapse/treated ascites samples harbored 31% (14–49), 59% (14– 124), and 90% (27–190) more mutations, neoantigens, and expressed neoantigens than primary/untreated ascites samples, respectively (*p* = 0.08,0.11,0.04 for the three tests). This trend was also apparent in a comparison of paired samples from the same donors (Figure S5).

**Figure 2:**
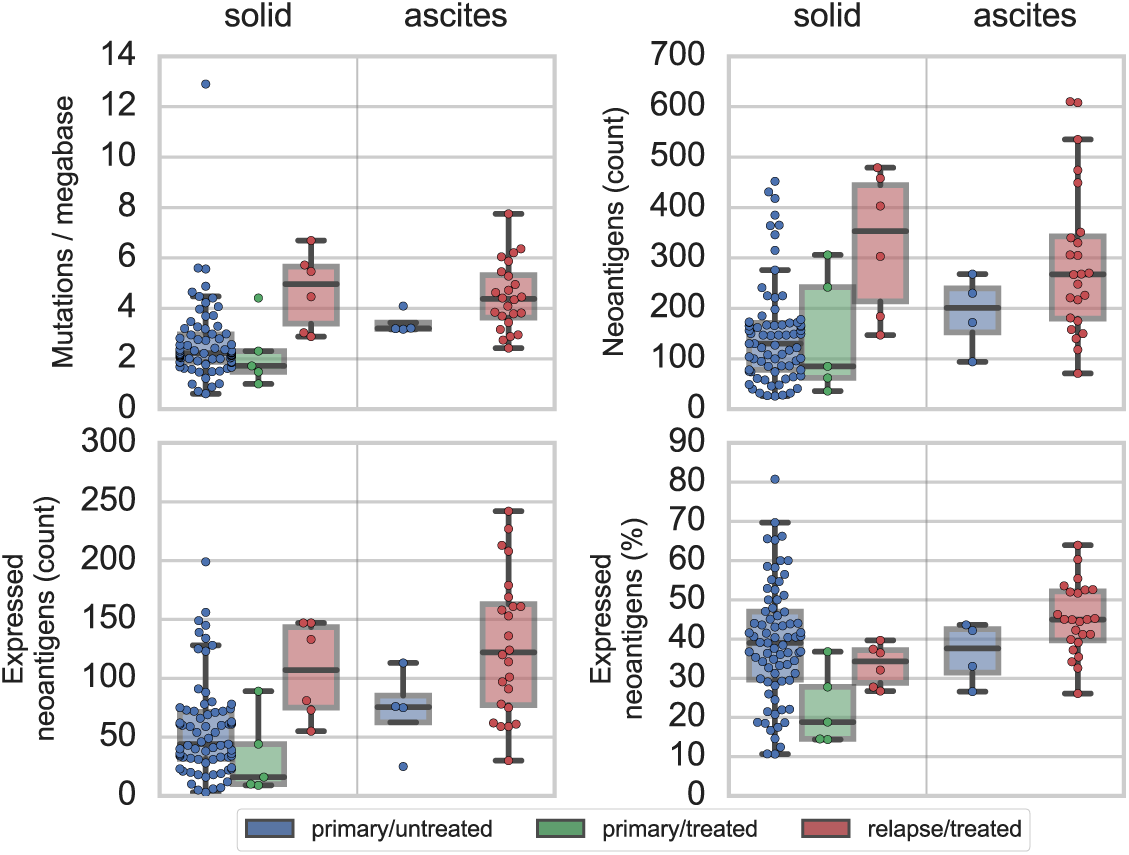
Stratified comparison of mutation and neoantigen burden of chemotherapy-treated and untreated samples. Mutations (upper left), neoantigens (upper right), and expressed neoantigens by count (lower left) and as a percent of total neoantigens (lower right) are shown for primary/untreated samples (blue; solid tumor n=75, ascites n=4), primary/treated samples (green; solid tumor n=5), and relapse/treated samples (red; solid tumor n=6, ascites n=24). The shaded boxes indicate the interquartile region and the median line. Points indicate individual samples.

In contrast, primary/treated samples, which were exposed to neoadjuvant chemotherapy (NACT) prior to surgery, did not exhibit increased numbers of mutations, neoantigens, or expressed neoantigens, and in fact trended toward decreased expressed neoantigen burden. The five primary/treated samples, all from solid tissue resections, harbored a median of 16 (9–89) expressed neoantigens compared to the median of 44 (39–60) observed in primary/untreated solid tissue samples, due to both fewer neoantigens in the DNA (median of 85 (36–306) vs. 130 (108–150)) and a lower rate of expression (median 19 (14–37) vs. 39 (36–42) percent of neoantigens). This trend did not reach significance (Mann-Whitney *p* = 0.09), and will require larger cohorts to assess.

### Chemotherapy signatures weakly contribute to neoantigen burden at relapse

While we cannot determine with certainty whether any particular mutation was chemotherapy-induced, we can estimate the fraction of mutations and neoantigens attributable to each signature in a sample (Figures 3 and S6).

**Figure 3:**
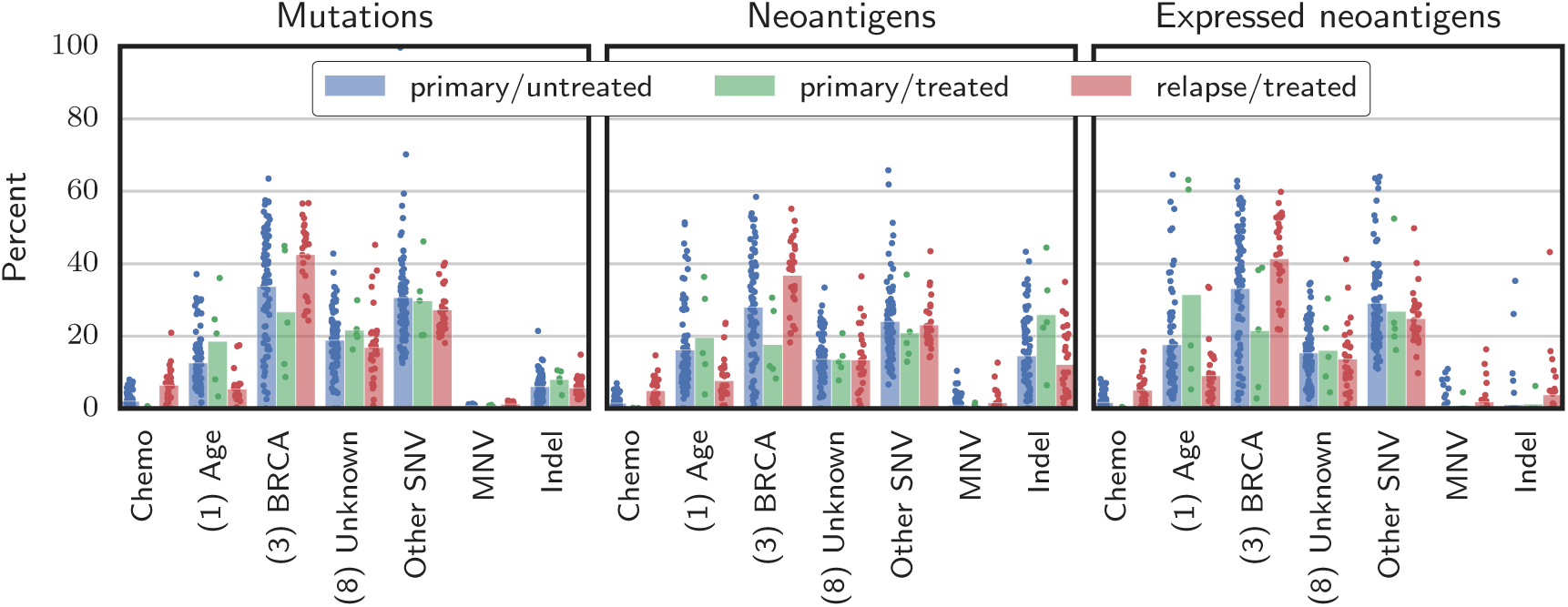
Contribution of key SNV signatures, MNVs, and indels on mutations *(left)*, neoantigens *(center)*, and expressed neoantigens *(right)*. The *Chemo* category combines the contributions from the chemotherapy signatures (cisplatin, cyclophosphamide, and etoposide). COSMIC signature numbers are in parentheses. The *Other SNV* category represents SNVs not accounted for by the signatures shown. Bars give the mean, and points indicate individual samples.

Similarly to results reported by Patch et al., the most prevalent mutational signatures in this cohort were COSMIC *Signature (3)*, associated with BRCA disruption, *Signature (8)*, of unknown etiology, and *Signature (1)*, associated with spontaneous deamination of 5-methylcytosine, a slow process active in healthy tissue that correlates with age (Figure S3 top and middle). These signatures together accounted for a median of 67% (95% CI 66–69) of mutations, 58% (56–61) of neoantigens, and 68% (67–71) expressed neoantigens across samples. These rates did not substantially differ with chemotherapy treatment.

The chemotherapy signatures accounted for a small but detectable part of the increased neoantigen burden of relapse samples. In primary/untreated samples, which indicate the background rate of chance attribution, chemotherapy mutational signatures accounted for a mean of 2% of the mutations (range 0–8), 2% (0–7) of the neoantigens, and 2% (0–8) of the expressed neoantigens. In each of the five primary/treated samples, less than 1% of the mutation, neoantigen, and expressed neoantigen burdens were attributed to chemotherapy signatures. For the relapse/treated samples, chemotherapy signatures accounted for a mean of 6% (range 0–21) of the mutations, 5% (0–15) of the neoantigens, and 5% (0–16) of the expressed neoantigens. The highest attribution to chemotherapy signatures occurred in sample AOCS-092-3-3, a relapse/treated sample from a patient who received five lines of platinum chemotherapy and eight distinct chemotherapeutic agents, the most in the cohort. For this sample, 21% (or approximately 3,200 of 15,491) of the SNVs, 15% (9 of 61) of the neoantigens, and 16% (5 of 30) of the expressed neoantigens were attributed to chemotherapy signatures.

Signature deconvolution considers only SNVs, but studies of platinum-induced mutations have also reported increases in the rate of dinucleotide variants and indels. Indeed, we observed more MNVs overall and specifically the platinum-associated MNVs *CT* → *AC* and *CA* → *AC* reported by Meier et al. [21] in treated patients in both absolute count and as a fraction of mutational burden (*p* < 10^*−6*^ for all tests). Sample AOCS-092-3-3, previously found to have the most chemotherapy-signature SNVs, also had the most platinum-associated dinucleotide variants and the second-most MNVs overall. This sample harbored 59 *CT → AC* or *CA → AC* mutations, compared to a mean of 3.2 (2.2–4.4) across all samples. Treated samples also harbored more indels in terms of absolute count (*p* = 10^−4^). Overall, while MNVs and indels generate more neoantigens per mutation than SNVs, they are rare, comprising less than 3% of the mutational burden and 13% of the neantigens in every sample (Figure 3), making it unlikely that chemotherapy-induced MNVs and indels have a large impact on neoantigen burden.

## Discussion

In this analysis of neoantigens predicted from DNA and RNA sequencing of ovarian cancer tumors and ascites samples, relapse samples obtained after chemotherapy exposure had a median of 90% more expressed neoantigens than untreated primary samples. However, our proposed chemotherapy mutational signatures accounted for no more than 16% of the expressed neoantigen burden in any sample. Most of the increase was instead attributable to mutagenic processes already at work in the primary samples, including COSMIC *Signature (3) BRCA* and *Signature (8) Unknown etiology*. Our results are in contrast to a study of NACT temozlomide-treated glioma, in which it was reported that over 98% of mutations detectable with bulk sequencing in some samples were attributable to temozolomide [5]. Whether this difference is due to the drug used or disease biology requires further study.

Detection of the cyclophosphamide and cisplatin signatures from the *G. Gallus* experiments showed some correlation with clinical treatment, whereas the *G. Gallus* etoposide and *C. Elegans* cisplatin signatures were not detected in chemotherapy-exposed samples. Many treated samples showed no chemotherapy signatures; when chemotherapy signatures were detected, they were found at levels close to the 6% detection threshold. In the case of cyclophosphamide, the deconvolution of all mutations from all samples identified the signature in 4/10 samples treated with cyclophosphamide and 4/104 unexposed samples. However, when we focused on mutations detected uniquely in the post-treatment paired samples, 6/8 samples exposed only to non-cyclophosphamide chemotherapies exhibited the signature. As it was rarely detected in pre-treatment samples, we suggest that the cyclophosphamide signature present in these post-treatment samples may reflect the effect of other chemotherapy, such as carboplatin, paclitaxel, doxorubicin, or gemcitabine. Analysis of the paired pre- and post-treatment samples indicated that the *G. Gallus* cisplatin signature was specific for cisplatin rather than carboplatin exposure, suggesting that carboplatin may induce fewer mutations or mutations with a different signature than cisplatin. The *C. Elegans* cisplatin signature may be less accurate than the *G. Gallus* cisplatin signature because it was derived from fewer mutations (784 vs. 2633) and from experiments of *C. Elegans* in various knockout backgrounds, which may not be relevant to these clinical samples. While only SNVs are accounted for by mutational signatures, an increase in indels and cisplatin-associated dinucleotide variants was observed in relapse/treated samples, but these variants remained relatively rare and generated less than 13% of the predicted neoantigen burden in every sample. Etoposide-induced mutations may be difficult to detect because in the *G. Gallus* experiments they occurred at a more uniform distribution of mutational contexts and at a much lower overall rate than mutations induced by cisplatin or cyclophosphamide. Importantly, only one patient in this cohort received etoposide.

The observed association between mutational signatures and clinical exposures gives some confidence that our analysis captures the effect of chemotherapy, but, as the preclinical signatures may differ from actual effects in patients, chemotherapy-induced mutations could be erroneously attributed to non-chemotherapy signatures. This would result in an underestimation of the impact of chemotherapy. We note, however, that the signatures dominant in the primary/untreated samples — COSMIC Signatures (1), (3), and (8) — also account for most of the SNVs in the relapse/treated samples. Therefore, irrespective of the accuracy of the chemotherapy signatures, it appears that most mutations in relapse samples are due to mutagenic processes operative prior to therapy.

NACT-treated tumors, which were exposed to chemotherapy as large tumors and for a short duration (typically 3 cycles), did not show increased mutation or neoantigen burden over untreated samples and had very few mutations attributed to chemotherapy. This is likely because neoadjuvant chemotherapy-induced mutations remain at undetectable allelic fractions without the population bottleneck created by surgery and/or the multiple lines of chemotherapy provided in the adjuvant setting.

We predicted a median of 64 (50–75) expressed MHC I neoantigens across all samples in the cohort, significantly more than the median of 6 recently reported by Martin et al. in this disease [14]. However, Martin et al. did not consider indels, MNVs, or multiple neoantigens that can result from the same missense mutation, used a 100nm instead of 500nm MHC I binding threshold, used predominantly lower quality (50bp) sequencing, and only explicitly considered HLA-A alleles. Predicted neoantigen burden is best considered a relative measure of tumor foreignness, not an absolute quantity readily comparable across studies.

This study has several important limitations. As it is based on bulk DNA sequencing of heterogeneous clinical samples, the analysis is limited to neoantigens arising from mutations that are present in at least 5-10% of the cells in a sample. Data from Patch et al. suggests that even late-stage disease remains polyclonal, therefore potentially obscuring the impact of chemotherapy on the tumor genome. While we may have been unable to detect subclonal mutations due to the depth of whole genome sequencing, it is expected that such clones would be unable to trigger an anti-tumor immune response that is effective against the bulk of the tumor [26]. Additionally, while the number of mutations attributed to signatures other than chemotherapy and those active in the primaries (COSMIC Signatures 1, 3, and 8) suggest that the preclinical signatures capture most chemotherapy-induced mutations, this reasoning assumes that chemotherapy does not induce mutations that are erroneously attributed to COSMIC Signatures 1, 3, or 8. Experiments using human cell lines exposed to the range of chemotherapies used in recurrent ovarian cancer may be needed to fully address this question. A further limitation is that this study does not consider neoantigens resulting from structural rearrangements such as gene fusions. Finally, this study relies on only 35 post-chemotherapy samples.

## Conclusion

In this study, we demonstrate a method for connecting mutational signatures extracted from studies of mutagen exposure in preclinical models with computationally predicted neoantigen burden in clinical samples. We found that relapsed high grade serous ovarian cancer tumors harbor nearly double the predicted expressed neoantigen burden of primary samples, and that cisplatin and cyclophophamide chemotherapy treatments account for a small but detectable part of this effect. The mutagenic processes responsible for most mutations at relapse are similar to those operative in primary tumors, with COSMIC *Signature (3) BRCA*, *Signature (1) Age*, and *Signature (8) Unknown etiology* accounting for most mutations and predicted neoantigens both before and after chemotherapy.

## 1 List of abbreviations

AOCS: Australian Ovarian Cancer Study, COSMIC: the Catalogue Of Somatic Mutations In Cancer, HGSC: high grade serous ovarian carcinoma, indel: an insertion or deletion mutation, MNV: multi nucleotide variant, NACT: neoadjuvant chemotherapy, SNV: single nucleotide variant

## 2 Ethics approval and consent to participate

The AOCS obtained ethics board approval at all institutions for patient recruitment, sample collection and research studies. Written informed consent was obtained from all participants in this study.

## 3 Consent for publication

Not applicable.

## 4 Availability of data and materials

All data generated during this study are included in this published article and its supplementary information files. The notebooks used to perform the analyses are available at https://github.com/hammerlab/paper-aocs-chemo-neoantigens.

## 7 Authors’ contributions

AS, DB, JH, and TO conceived and coordinated the study. TO performed the research and wrote the manuscript. EC curated the clinical records. AA, BAA, and JB advised on analysis methods. All authors revised the manuscript critically.

## 8 Acknowledgements

We thank Leonid Rozenberg for assistance with sequence-based HLA typing. We also thank Dariush Etemadmoghadam and Ann-Marie Patch at Peter MacCallum Cancer Centre for assistance accessing AOCS data sets.

## Additional Files

**Figure S1:**
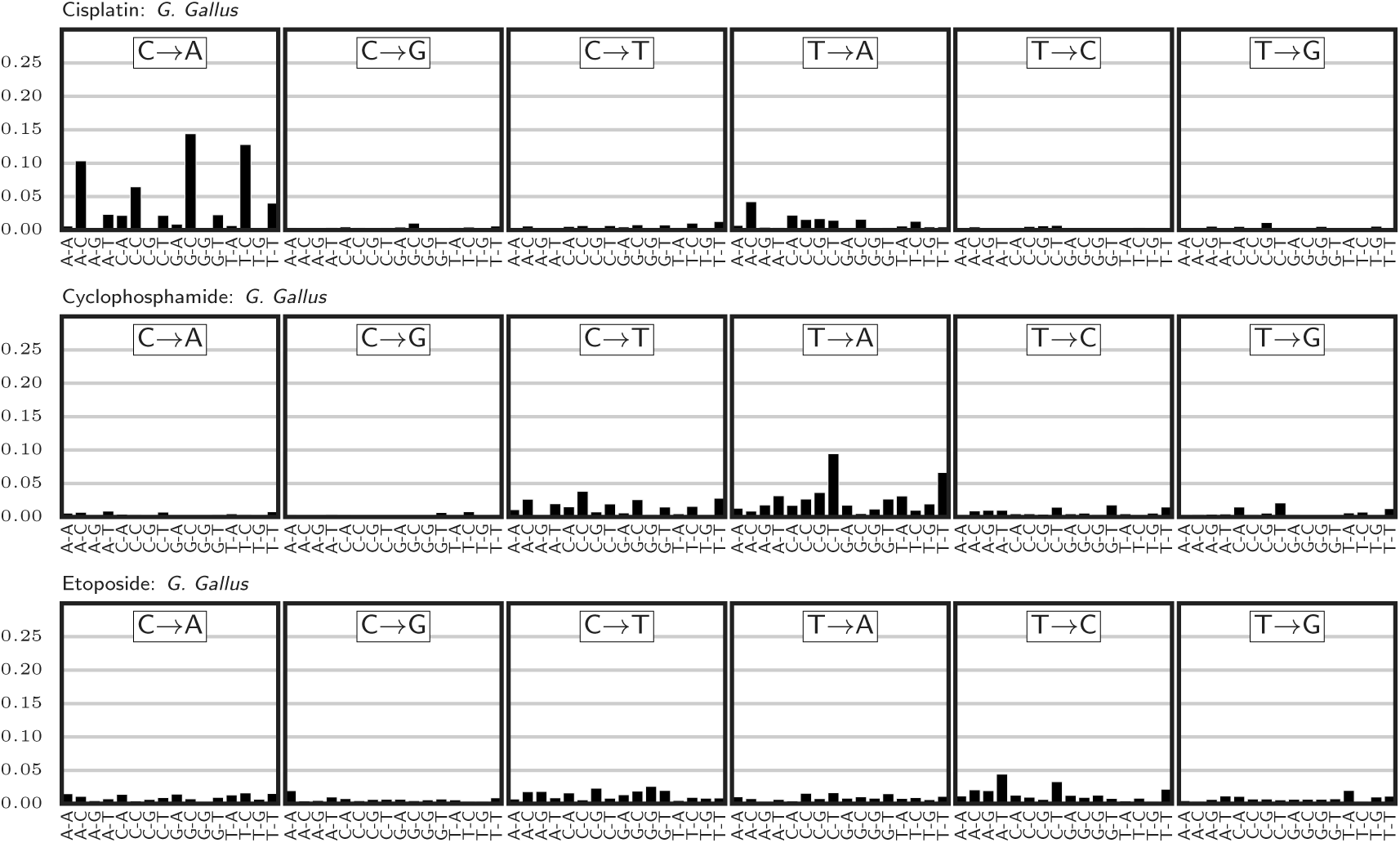
Mutational signatures extracted from Szikriszt et al. [20]

Additional Files
| Data File | Format | Title | Description |
| --- | --- | --- | --- |
| Additional File 1.csv | CSV | Samples | Sample identifiers, basic clinical information, specimen purities, mutation and neoantigen burden, contributions of major mutational signatures to mutations and neoantigens, and chemotherapy treatments |
| Additional File 2.csv.zip | CSV zip | Mutations | Mutations, read counts, predicted effects, and resulting neoantigens |
| Additional File 3.csv | CSV | HLA types | Donor HLA types |
| Additional File 4.csv | CSV | Mutational signatures | COSMIC signatures and extracted chemotherapy signatures |
| Additional File 5.csv | CSV | Signature deconvolutions | Results of mutational signature deconvolution, including separate analyses of all mutations and mutations unique to the treated paired samples |
| Additional File 6.csv | CSV | Shared neoantigens | Neoantigens predicted for multiple patients |

**Figure S2:**
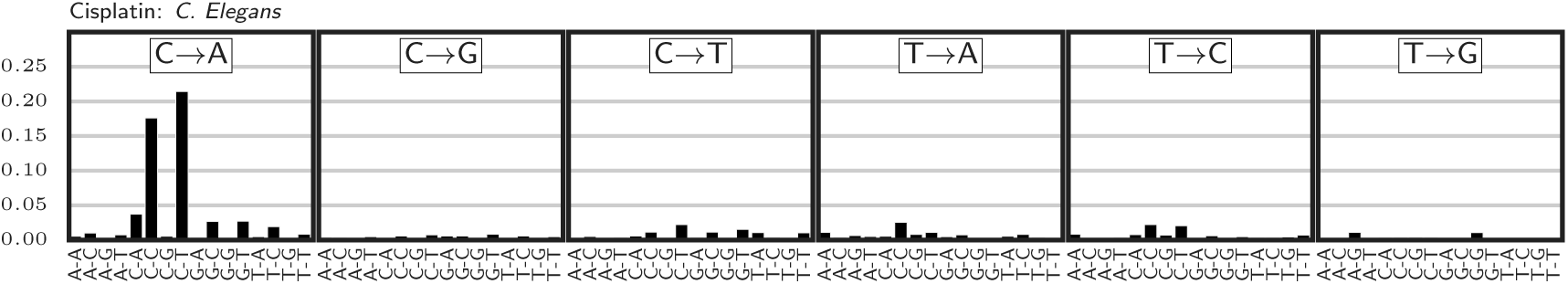
Mutational signature extracted from Meier et al. [21]

**Figure S3:**
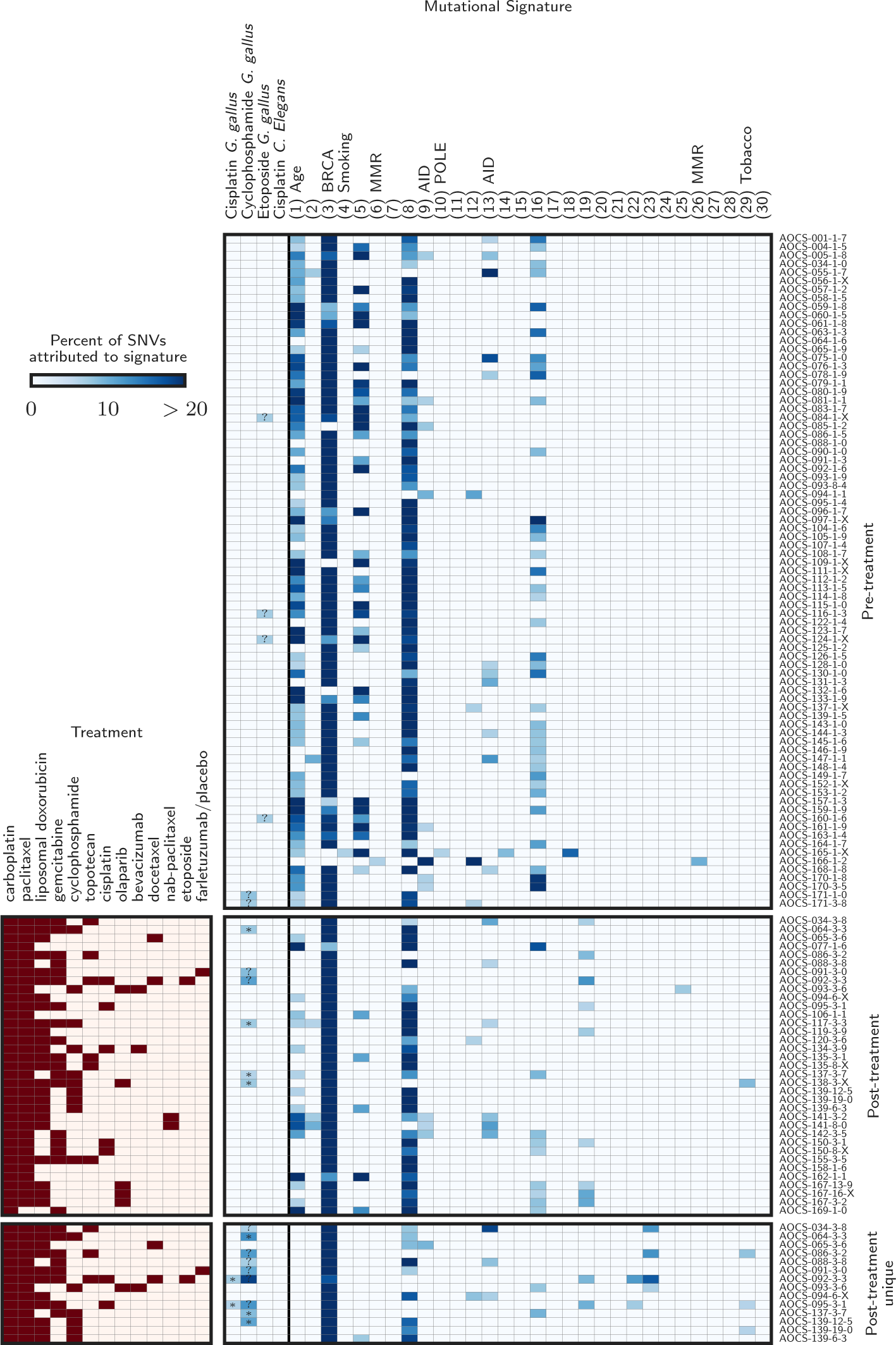
Detected mutational signatures across all samples. The symbols are as in main text Figure 1. The top and middle panels show the signature deconvolutions for all pre- and post-treatment samples, respectively. The bottom panel shows deconvolutions for the mutations unique to the paired post-treatment samples, requiring high coverage and no variant reads in the donor-matched pre-treatment sample.

**Figure S4:**
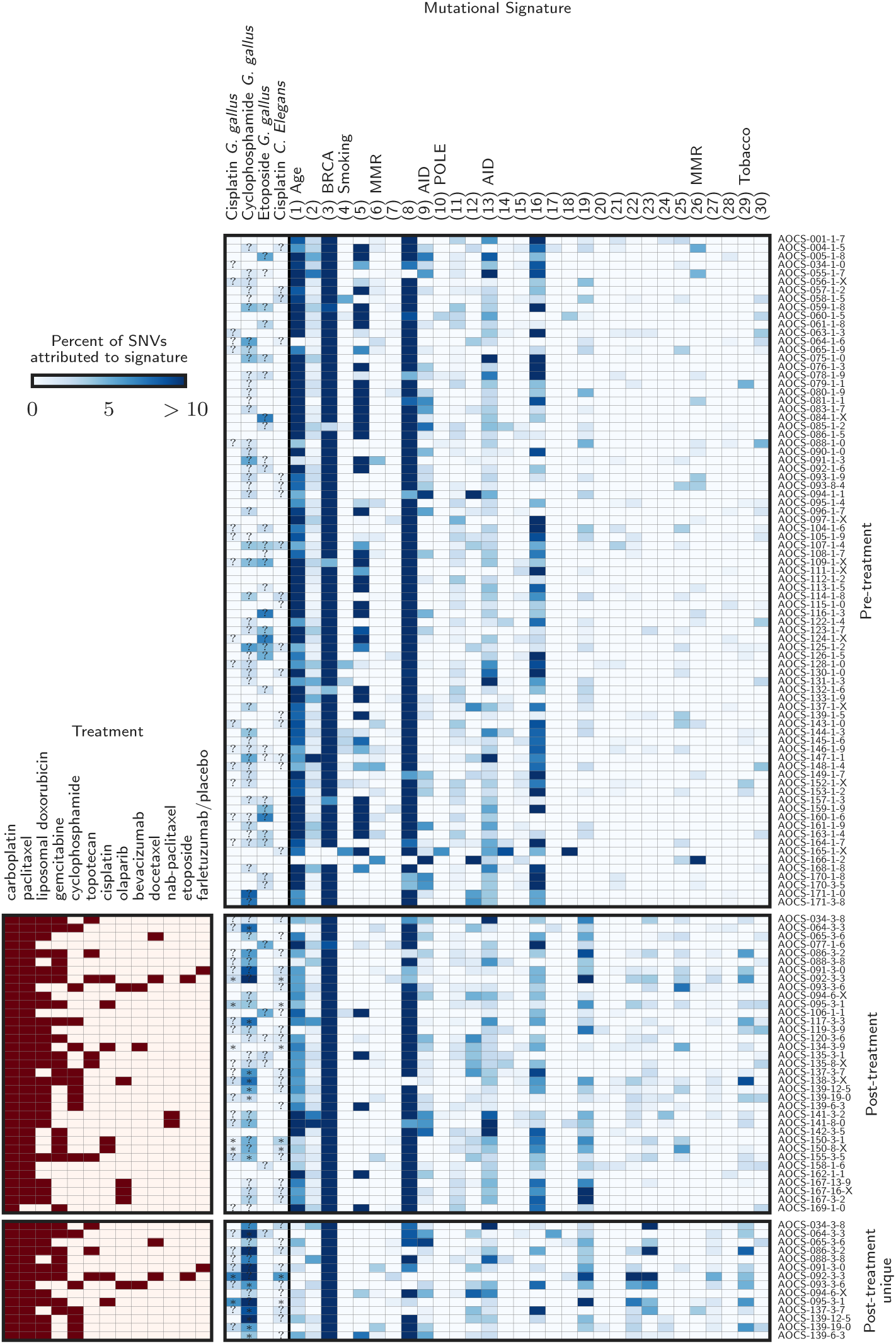
Mutational signature deconvolutions without any threshold of detection. Here, signatures accounting for less than the 6% recommended detection threshold are included. See Figure S3.

**Figure S5:**
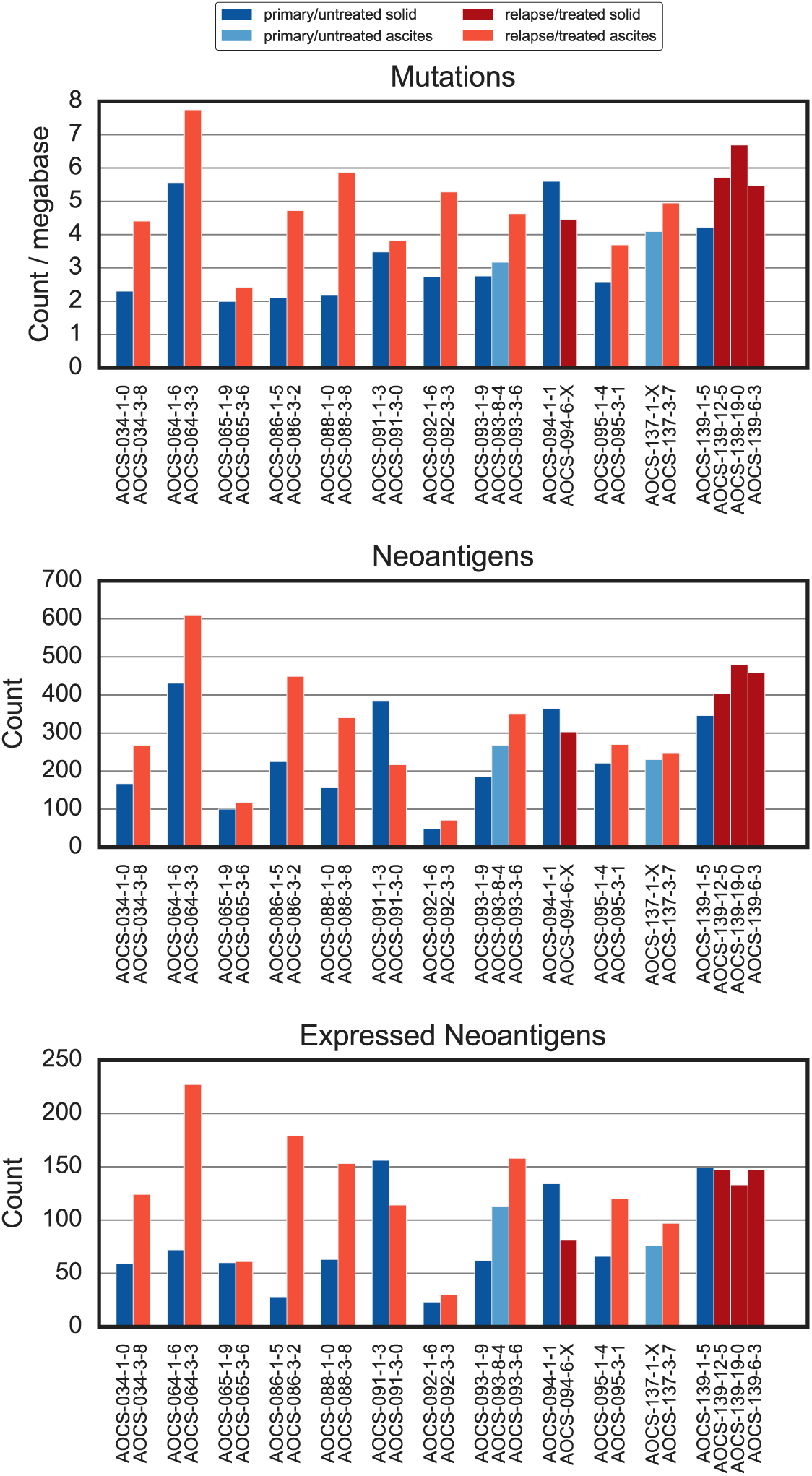
Mutations, neoantigens, and expressed neoantigens for donor-matched primary/untreated and relapse/treated samples.

**Figure S6:**
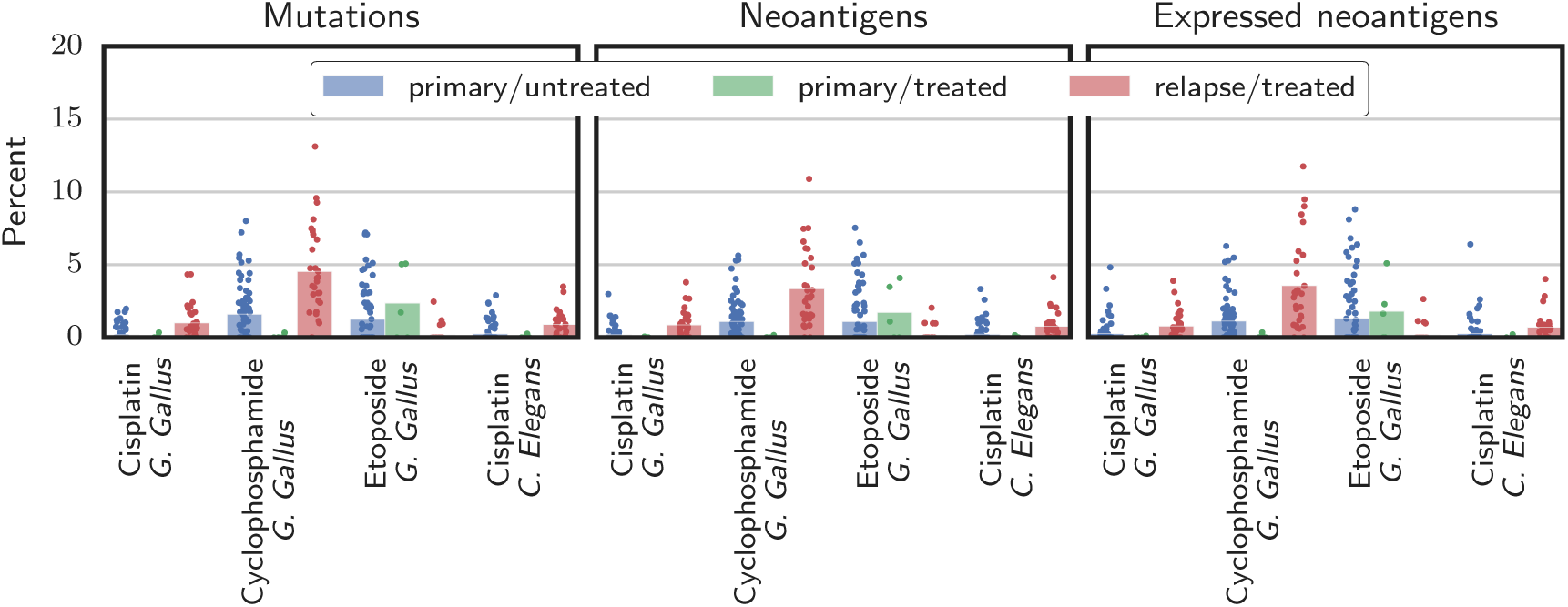
Contribution of chemotherapy SNV signatures. The fraction of each sample’s mutations, neoantigens, and expressed neoantigens attributed to putative chemotherapy signatures is shown. Bars give the mean, and points indicate individual samples.

## References

1. Hannan MA, Al-Dakan AA, Hussain SS, Amer MH. Mutagenicity of cisplatin and carboplatin used alone and in combination with four other anticancer drugs. Toxicology [Internet]. Elsevier BV; 1989;55:183–91. Retrieved from: http://dx.doi.org/10.1016/0300-483x(89)90185-6

2. Anderson D, Bishop JB, Garner RC, Ostrosky-Wegman P, Selby PB. Cyclophosphamide: Review of its mutagenicity for an assessment of potential germ cell risks. Mutation Research/Fundamental and Molecular Mechanisms of Mutagenesis [Internet]. Elsevier BV; 1995;330:115–81. Retrieved from: http://dx.doi.org/10.1016/0027-5107(95)00039-l

3. Nakanomyo H, Hiraoka M, Shiraya M. Mutagenicity tests of etoposide and teniposide. J. Toxicol. Sci. [Internet]. Japanese Society of Toxicology; 1986;11:301–10. Retrieved from: http://dx.doi.org/10.2131/jts.11.supplementi301

4. Ding L, Ley TJ, Larson DE, Miller CA, Koboldt DC, Welch JS, et al. Clonal evolution in relapsed acute myeloid leukaemia revealed by whole-genome sequencing. Nature [Internet]. Nature Publishing Group; 2012;481:506–10. Retrieved from: http://dx.doi.org/10.1038/nature10738

5. Johnson BE, Mazor T, Hong C, Barnes M, Aihara K, McLean CY, et al. Mutational Analysis Reveals the Origin and Therapy-Driven Evolution of Recurrent Glioma. Science [Internet]. American Association for the Advancement of Science (AAAS); 2013;343:189–93. Retrieved from: http://dx.doi.org/10.1126/science.1239947

6. Murugaesu N, Wilson GA, Birkbak NJ, Watkins TBK, McGranahan N, Kumar S, et al.Tracking the Genomic Evolution of Esophageal Adenocarcinoma through Neoadjuvant Chemotherapy.Cancer Discovery [Internet]. American Association for Cancer Research (AACR); 2015;5:821–31. Retrieved from: http://dx.doi.org/10.1158/2159-8290.cd-15-0412

7. .Chen DS, Mellman I. Oncology Meets Immunology: The Cancer-Immunity Cycle. Immunity [Internet]. Elsevier BV; 2013;39:1–10. Retrieved from: http://dx.doi.org/10.1016/j.immuni.2013.07.012

8. Schumacher TN, Schreiber RD. Neoantigens in cancer immunotherapy. Science [Internet]. American Association for the Advancement of Science (AAAS); 2015;348:69–74. Retrieved from: http://dx.doi.org/10.1126/science.aaa4

9. Lundegaard C, Lund O, Kesmir C, Brunak S, Nielsen M. Modeling the adaptive immune system: predictions and simulations. Bioinformatics [Internet]. Oxford University Press (OUP); 2007;23:3265–75. Retrieved from: http://dx.doi.org/10.1093/bioinformatics/btm471

10. Allen EMV, Miao D, Schilling B, Shukla SA, Blank C, Zimmer L, et al. Genomic correlates of response to CTLA-4 blockade in metastatic melanoma. Science [Internet]. American Association for the Advancement of Science (AAAS); 2015;350:207–11. Retrieved from: http://dx.doi.org/10.1126/science.aad0095

11. Rizvi NA, Hellmann MD, Snyder A, Kvistborg P, Makarov V, Havel JJ, et al. Mutational landscape determines sensitivity to PD-1 blockade in non-small cell lung cancer. Science [Internet]. American Association for the Advancement of Science (AAAS); 2015;348:124–8. Retrieved from: http://dx.doi.org/10.1126/science.aaa1348

12. Lawrence MS, Stojanov P, Polak P, Kryukov GV, Cibulskis K, Sivachenko A, et al.Mutational heterogeneity in cancer and the search for new cancer-associated genes. Nature [Internet]. Springer Nature; 2013;499:214–8. Retrieved from: http://dx.doi.org/10.1038/nature12213

13. Patch A-M, Christie EL, Etemadmoghadam D, Garsed DW, George J, Fereday S, et al. Whole—genome characterization of chemoresistant ovarian cancer. Nature [Internet]. Nature Publishing Group; 2015;521:489–94. Retrieved from: http://dx.doi.org/10.1038/nature14410

14. Martin SD, Brown SD, Wick DA, Nielsen JS, Kroeger DR, Twumasi-Boateng K, et al. Low Mutation Burden in Ovarian Cancer May Limit the Utility of Neoantigen-Targeted Vaccines. Ashkar A A, editor. PLOS ONE [Internet]. Public Library of Science (PLoS); 2016;11:e0155189.Retrieved from: http://dx.doi.org/10.1371/journal.pone.0155189

15. Lab H. Varcode [Internet]. Varcode GitHub. Hammer Lab; 2016. Retrieved from: https://github.com/hammerlab/var

16. Boegel S, L¨ower M, Sch¨afer M, Bukur T, Graaf J de, Boisgu´erin V, et al. HLA typing from RNA-Seq sequence reads. Genome Medicine [Internet]. Springer Science Business Media; 2012;4:102. Retrieved from: http://dx.doi.org/10.1186/gm403

17. Szolek A, Schubert B, Mohr C, Sturm M, Feldhahn M, Kohlbacher O. OptiType: precision HLA typing from next-generation sequencing data. Bioinformatics [Internet]. Oxford University Press (OUP); 2014;30:3310–6. Retrieved from: http://dx.doi.org/10.1093/bioinformatics/btu548

18. Lundegaard C, Lamberth K, Harndahl M, Buus S, Lund O, Nielsen M. NetMHC-3.0:accurate web accessible predictions of human mouse and monkey MHC class I affinities for peptides of length 8-11. Nucleic Acids Research [Internet]. Oxford University Press (OUP); 2008;36:W509–W512. Retrieved from: http://dx.doi.org/10.1093/nar/gkn202

19. Alexandrov LB, Nik-Zainal S, Wedge DC, Aparicio Sa JR, Behjati S, Biankin AV, et al. Signatures of mutational processes in human cancer. Nature [Internet]. 2013; 500: 415–21. Retrieved from: http://www.pubmedcentral.nih.gov/articlerender.fcgi?artid=3776390&tool=pmcentrez&rendertype=abstract

20. Rosenthal R, McGranahan N, Herrero J, Taylor BS, Swanton C. deconstructSigs: delineating mutational processes in single tumors distinguishes DNA repair deficiencies and patterns of carcinoma evolution. Genome Biol [Internet]. Springer Science Business Media; 2016; 17. Retrieved from: http://dx.doi.org/10.1186/s13059-016-0893-4

21. Meier B, Cooke SL, Weiss J, Bailly AP, Alexandrov LB, Marshall J, et al. C. elegans whole-genome sequencing reveals mutational signatures related to carcinogens and DNA repair deficiency. Genome Research [Internet]. Cold Spring Harbor Laboratory Press; 2014; 24: 1624–36. Retrieved from: http://dx.doi.org/10.1101/gr.175547.114

22. Szikriszt B, P´oti Ad´am, Pipek O, Krzystanek M, Kanu N, Moln´ar J, et al. A comprehensive survey of the mutagenic impact of common cancer cytotoxics. Genome Biol [Internet]. Springer Science Business Media; 2016;17. Retrieved from: http://dx.doi.org/10.1186/s13059-016-0963-7

23. Gelman A, Lee D, Guo J. Stan: A Probabilistic Programming Language for Bayesian Inference and Optimization. Journal of Educational and Behavioral Statistics [Internet]. American Educational Research Association (AERA); 2015; 40:530–43. Retrieved from: http://dx.doi.org/10.3102/1076998615606113

24. Institute WTS. Signatures of Mutational Processes in Human Cancer [Internet]. http://cancer.sanger.ac.uk/cosmic/signatures; 2016. Retrieved from: http://cancer.sanger.ac.uk/cosmic/signatures

25. Sette A, Vitiello A, Reherman B, Fowler P, Nayersina R, Kast WM, et al. The relationship between class I binding affinity and immunogenicity of potential cytotoxic T cell epitopes. Journal of immunology (Baltimore, Md.: 1950) [Internet]. 1994;153:5586–92. Retrieved from: http://www.ncbi.nlm.nih.gov/pubmed/7527444

26. McGranahan N, Furness AJS, Rosenthal R, Ramskov S, Lyngaa R, Saini SK, et al. Clonal neoantigens elicit T cell immunoreactivity and sensitivity to immune checkpoint blockade. Science [Internet]. American Association for the Advancement of Science (AAAS); 2016;351:1463–9. Retrieved from: http://dx.doi.org/10.1126/science.aaf1490

27. Dunn GP, Bruce AT, Ikeda H, Old LJ, Schreiber RD. Cancer immunoediting: from immunosurveillance to tumor escape. Nature Immunology [Internet]. Nature Publishing Group; 2002;3:991–8. Retrieved from: http://dx.doi.org/10.1038/ni1102-991

28. Demaria S, Volm MD, Shapiro RL, Yee HT, Oratz R, Formenti SC, et al. Development of Tumor-infiltrating Lymphocytes in Breast Cancer after Neoadjuvant Paclitaxel Chemotherapy. 2001;7:3025–30.

29. Wu X, Feng Q-M, Wang Y, Shi J, Ge H-L, Di W. The immunologic aspects in advanced ovarian cancer patients treated with paclitaxel and carboplatin chemotherapy. Cancer Immunology Immunotherapy [Internet]. Springer Science Business Media; 2009;59:279–91. Retrieved from: http://dx.doi.org/10.1007/s00262-009-0749-9

30. Pfannenstiel LW, Lam SSK, Emens LA, Jaffee EM, Armstrong TD. Paclitaxel enhances early dendritic cell maturation and function through TLR4 signaling in mice. Cellular Immunology [Internet]. Elsevier BV; 2010;263:79–87. Retrieved from: http://dx.doi.org/10.1016/j.cellimm.2010.03.001

31. Hodge JW, Garnett CT, Farsaci B, Palena C, Tsang K-Y, Ferrone S, et al. Chemotherapy-induced immunogenic modulation of tumor cells enhances killing by cytotoxic T lymphocytes and is distinct from immunogenic cell death. International Journal of Cancer [Internet]. Wiley-Blackwell; 2013;133:624–36. Retrieved from: http://dx.doi.org/10.1002/ijc.28070

